# EEG resting state alpha dynamics predict individual proneness to auditory hallucinations

**DOI:** 10.1101/2023.05.22.541696

**Authors:** H. Honcamp, S.X. Duggirala, J. Rodino Climent, A. Astudillo, N.J. Trujillo-Barreto, M. Schwartze, D.E.J. Linden, T.A.M.J. van Amelsvoort, W. El-Deredy, S.A. Kotz

## Abstract

**Introduction:** Auditory verbal hallucinations (AVH) are a transdiagnostic phenomenon but also occur in the general population. The disposition to experience AVH is considered a continuous expression from non-clinical to clinical hallucination proneness (HP). Currently, little is known about the neurophysiology of the non-clinical HP part of the continuum. AVH might result from a heightened sensitivity to sensory inputs and a decreased ability to differentiate between externally and internally generated input. Resting state (RS) alpha band activity is associated with perceptual sensitivity, attentional shifts, and cognitive control. Accordingly, spontaneous alpha fluctuations might present as a HP correlate. To investigate the time-varying dynamics of alpha band activity, we deployed a novel method for brain state allocation.

**Methods:** We recorded RS electroencephalography (EEG) data from 33 individuals with varying levels of HP but without clinically relevant hallucinations and used a Hidden Semi-Markov Model (HsMM) to identify five recurrent alpha states with unique temporal dynamics and topographies. The states’ mean duration and occupancy were analyzed as a function of HP. The sources of each state were reconstructed to identify the most active brain areas and their correspondence with known resting state networks.

**Results:** Occupancy and mean duration of a state corresponding to sensorimotor, auditory, and default-mode network (DMN) areas significantly predicted auditory and auditory-verbal HP, but not general HP. The temporal dynamics of all other states did not relate to HP.

**Conclusion:** Alpha brain state sources align with prior results on the role of the alpha in the DMN. The temporal dynamics of alpha might reflect individual differences for attentional biases to internally generated sensory events and altered auditory perceptual sensitivity. Thus, changes in the temporal brain state dynamics of RS alpha oscillations could present as a neural marker of increased vulnerability to auditory hallucinatory experiences.

## Introduction

### Auditory verbal hallucinations and hallucination proneness

Auditory verbal hallucinations (AVH) are a cardinal symptom of schizophrenia, but also occur in other neuropsychiatric disorders with psychotic features and the general population (Bartels-Velthuis, Jenner, van de Willige, van Os, & Wiersma, 2010; Linszen et al., 2022). Thus, AVH are not necessarily related to need for care but rather lie on a severity continuum, referred to as hallucination proneness (HP) (Johns et al., 2014). An increased frequency of occurrence and severity of AVH is associated with a higher risk of developing psychotic symptoms captured by the psychosis-continuum hypothesis (Johns & Van Os, 2001). The continuum perspective suggests that the underlying cognitive mechanisms and neuronal changes associated with AVH in healthy individuals are an attenuated version of those observed in clinically diagnosed individuals (e.g., schizophrenia patients). Correspondingly, individuals on the upper end of the non-clinical HP continuum might display a heightened vulnerability to transition to clinical AVH. This continuum perspective is empirically supported by various behavioral, electrophysiological, structural, and functional neuroimaging studies (Allen et al., 2005; Badcock & Hugdahl, 2012; Diederen et al., 2012; Marschall et al., 2023; Paulik, Badcock, & Maybery, 2008; Pinheiro, Farinha-Fernandes, Roberto, & Kotz, 2019).

### Auditory verbal hallucinations and the brain’s resting state

AVH seem to result from altered inner speech monitoring, aberrant salience and source misattribution, and changes in cognitive control mechanisms, e.g., the reality checking of verbal intrusions from long-term memory (Badcock & Hugdahl, 2012). In other words, voice hearers fail to differentiate between internally and externally generated sensory events. Corresponding neural correlates of AVH include event-related potential changes indicative of aberrant sensory feedback to voices, altered functional connectivity (FC) in language and auditory brain regions, and functional interactions between respective cortical networks (Diederen et al., 2012; Pinheiro et al., 2020; Spray, Beer, Bentall, Sluming, & Meyer, 2018). However, as most research has focused on clinical and non-clinical voice-hearing, little is known about the underlying neurophysiology of the non-clinical HP continuum where symptoms are not (yet) present. Considering the fluctuating and unprovoked nature of AVH, the temporal dynamics of the resting brain might be a suitable target to elucidate their neural basis (Alderson-Day, McCarthy-Jones, & Fernyhough, 2015; Northoff & Qin, 2011). The brain’s resting state (RS) transitions between patterns of intrinsically and spontaneously generated neural activity and is associated with increased engagement of so-called resting state networks (RSNs). RSNs consist of functionally relevant cortical, subcortical, and cerebellar brain regions, interacting systematically in time. The FC patterns resulting from the network’s systematic (dis-)engagement inform about healthy and pathological brain function (Damoiseaux et al., 2006). The ‘resting state hypothesis of AVH’ proposed by Northoff and Qin (2011) suggests that AVH result from a dysfunctional interaction, i.e., aberrant bottom-up and top-down processes between the Default-Mode Network (DMN) and other RSNs, including the salience, cognitive control, and auditory networks. This is in line with the predictive coding framework of AVH, suggesting that voice hearing results from an impairment in predicting (and updating predictions of) sensory information. Correct encoding and prediction of sensory information induces an attenuation of neural activity in the respective sensory networks which minimizes prediction errors and facilitates the differentiation between internally and externally generated stimuli. Disruptions in the prediction-based neural suppression thus promote the perception of voices that are not actually present (Horga, Schatz, Abi-Dargham, & Peterson, 2014).

### Resting state dynamics

The brain’s RS is inherently non-stationary and dynamic (Hutchison et al., 2013), i.e., it fluctuates between distinct FC patterns on a sub-second timescale (Baker et al., 2014; Trujillo-Barreto, Araya, & El-Deredy, 2019; Vidaurre, Smith, & Woolrich, 2017; Woolrich et al., 2013). Accordingly, RS time series can be segmented into a sequence of recurring states, characterized by unique activity profiles and temporal dynamics (Baker et al., 2014; Kottaram et al., 2019; Trujillo-Barreto et al., 2019; Vidaurre et al., 2018). However, these states are “hidden”, as they cannot be observed directly but need to be inferred (Trujillo-Barreto et al., 2019). To this end, different segmentation methods can be used, e.g., sliding window analysis, microstate analysis, and computational modeling approaches (Andreou et al., 2014; Geng et al., 2020; Kottaram et al., 2019). These approaches aim at characterizing neural time series as a sequence of recurrent connectivity states, as their order and switching dynamics relate to cognition and behavior in healthy and patient populations (Kottaram et al., 2019; Nishida et al., 2013; Vidaurre et al., 2016).

The sliding window method is commonly applied to functional magnetic resonance imaging (fMRI) data and provides high spatial accuracy when localizing dysfunctional brain processes. Applications of this method in clinical voice hearers emphasized a key role of DMN dysfunction in AVH (Geng et al., 2020; Weber et al., 2020). This result supports previous findings in clinical (Manoliu et al., 2014) and non-clinical voice hearers (van Lutterveld, Diederen, Otte, & Sommer, 2014) as well as healthy individuals with an increased psychosis risk (Hua et al., 2019). Together, these findings yield important evidence for altered RSN interactions associated with a predisposition to AVH. Another recently proposed method to investigate time varying AVH dynamics, called the leading eigenvector dynamics analysis (LEiDA), was applied to fMRI time series of clinical and non-clinical voice hearers (Marschall et al., 2023). The results showed that network dynamics differed substantially between pure rest and periods during which participants hallucinated but were similar in clinical and non-clinical AVH. However, fMRI only captures an indirect measure of neural activity and lacks the required temporal sensitivity to monitor the sub-second dynamics of the resting brain (Honcamp, Schwartze, Linden, El-Deredy, & Kotz, 2022; Hutchison et al., 2013). Moreover, the sliding window method is biased toward FC transitions that occur within predefined and fixed time windows and thus overlooks transitions that occur outside of those windows. We therefore require higher temporal resolution methods such as magneto-or electroencephalography (M/EEG) as well as tools that adequately capture temporal dynamics on all timescales (Hutchison et al., 2013; Trujillo-Barreto et al., 2019). Alternatively, the EEG microstate analysis assesses topographical changes at global field power (GFP) peaks across time to cluster RS time series into four classes of global scalp activity patterns, referred to as microstate classes A-D (Lehmann, Strik, Henggeler, König, & Koukkou, 1998; Michel & Koenig, 2018). This approach yielded robust evidence for altered microstate durations in persons with AVH, indicating altered information processing at rest (Andreou et al., 2014; Kindler, Hubl, Strik, Dierks, & Koenig, 2011; Nishida et al., 2013). However, the microstate approach primarily focuses on topographical changes at GFP peaks, and the switching dynamics are not considered during the analysis but are inferred a posteriori. Thus, the microstate analysis may miss subtle and short-lasting signatures of RSN interaction (Honcamp et al., 2022). We therefore currently lack a clear understanding of the dynamic brain changes that might explain the AVH phenomena holistically and along the HP continuum.

### Characterizing brain state dynamics through Hidden semi-Markov Modeling

Computational generative models such as the Hidden Markov Model (HMM) can be used to accurately characterize the brain’s non-stationarity in a data-driven manner and to draw conclusions about underlying temporal dynamics of the brain’s RS at a sub-second timescale (Baker et al., 2014). HMMs iteratively learn the statistical properties (e.g., means and covariances) of the data directly and characterize time series as a sequence of distinct and recurrent states that are defined by a quasi-stable activity patterns and distinct temporal properties. The state sequence, therefore, reveals relevant information about network connectivity and switching dynamics, i.e., how long a given state is active, how much time it occupies, and how likely it transitions to other states. HMMs have been successfully applied to M/EEG and fMRI data (Baker et al., 2014; Hunyadi, Woolrich, Quinn, Vidaurre, & De Vos, 2019; Trujillo-Barreto et al., 2019; Woolrich et al., 2013).

The Hidden Semi-Markov Model (HsMM) is a generalization of the HMM that allows incorporating more realistic assumptions about long-range dependencies of M/EEG time series and network switching behavior (Trujillo-Barreto et al., 2019). Correspondingly, the HsMM allows for explicit modeling state durations. As RSNs systematically alternate between periods of activation and deactivation, the states’ activity duration and occupancy may reflect stability of underlying FC signatures and thereby inform about (in-)efficient information processing. In contrast to the sliding window approach and the EEG microstate analysis, the HsMM learns the relevant timescales directly from the data and is not restricted to a predefined time window or topographical fluctuations at GFP peaks (Rukat, Baker, Quinn, & Woolrich, 2016; Trujillo-Barreto et al., 2019). Thus, the HsMM overcomes the main limitations of previously applied approaches and is considered an adequate alternative to reveal neural correlates of HP (Honcamp et al., 2022).

### The role of alpha in the brain’s resting state

Alpha band activity (8-12 Hz) is associated with cortical excitability through selective attention across sensory domains (Hindriks, Micheli, Mantini, & Deco, 2017). Specifically, alpha power fluctuations modulate the sensitivity to sensory experiences, i.e., they alter the threshold at which sensory stimuli are perceived, including visual, tactile, and pain stimuli (Craddock, Poliakoff, El-Deredy, Klepousniotou, & Lloyd, 2017; Ecsy, Jones, & Brown, 2017; Klimesch, Sauseng, & Hanslmayr, 2007). Moreover, alpha band activity can influence the temporal and spatial grouping of sensory information when perceiving ambiguous stimuli, indicating that alpha band activity shapes the way sensory information is attended to and perceived (Shen, Han, Chen, & Chen, 2019). Given that changes in alpha band activity are observed across multiple modalities, the same mechanisms could contribute to the detection of auditory ‘phantom’ percepts. In fact, alpha power modulations with a parieto-occipital topography in eyes-closed conditions synchronize with directing attention toward rather than away from auditory/speech stimuli (Tune, Wöstmann, & Obleser, 2018; Wöstmann, Schmitt, & Obleser, 2020). Alpha fluctuations are also associated with cognitive control such as the maintenance of cognitive representations (Clements et al., 2021). In turn, deficits in cognitive control such as intrusive thoughts and impaired reality-checking, are frequently related to HP, psychosis proneness, and schizotypy (Alderson-Day et al., 2019; Paulik et al., 2008; Waters et al., 2012). Although the canonical EEG frequency bands (delta, theta, alpha, beta, gamma) cannot be exclusively mapped onto one RSN, there is evidence that links alpha band activity to the DMN (Hillebrand, Barnes, Bosboom, Berendse, & Stam, 2012; Samogin et al., 2020). Given the sub-second fluctuations of RS activity, the dominant role of alpha band activity in ‘attended’ sensations, and the link to DMN activation, the analysis of RS alpha band dynamics might be particularly suited to reveal neural correlates along the HP continuum.

### Objectives and relevance

Currently, we lack evidence of altered brain state dynamics in hallucination-prone individuals on the non-clinical spectrum using high temporal resolution measures. The HsMM ensures the high temporal sensitivity necessary to characterize subtle changes in dynamic RS connectivity. It is therefore an adequate means to elucidate aberrant state dynamics that increase AVH vulnerability. Considering that alpha modulations of can reflect individual sensitivities to sensory experiences, the temporal dynamics of the brain’s RS in the alpha band might be particularly informative to uncover the neural underpinnings of non-clinical HP.

The current study assessed the temporal dynamics of RS EEG alpha activity as a function of HP. To this end, we applied an HsMM to describe the RS EEG alpha rhythm envelope of 33 individuals to analyze the predictive value of state durations (how long a state is active on average) and occupancy (how much time a state occupies relative to the total recording time) for general, auditory, and auditory-verbal HP. We hypothesized that the HsMM-derived brain state dynamics would show sub-second temporal dynamics and between-subject differences thereof. We further expected the alpha state dynamics to vary as a function of HP and thus be informative for eventual AVH risk assessment and theories of the psychosis continuum in general. Moreover, we aimed to identify the sources corresponding to each state by means of frequency-domain source localization. We expected alpha state brain sources to correspond to well-known RSNs, predominantly the DMN.

## Methods

### Participants and procedure

The study was part of a larger multi-modal research line, focusing on altered brain dynamics in non-clinical and clinical voice hearers. This research line was conducted at Maastricht University, the Netherlands. Ethical approval was granted by the medical ethics committee responsible (METC-20 035; the study was prematurely terminated due to recruitment difficulties). Inclusion criteria for participation were age between 16-65 years, normal or corrected hearing, no neurological disorder (e.g., epilepsy, tumor, lesion), and no previous head or brain injury. We collected RS EEG data of 38 participants from the general population. Based on standard data quality assessment, thirty-three individuals (9 males, 24 females; Mean age = 23,36; SD = 2.65) were selected for the current study. The sample consisted of university students (one recent graduate), whose participation was either rewarded monetarily with vouchers or study credits. All participants provided written informed consent before participation.

### Data acquisition

Participants were invited for two sessions, including a neuropsychological assessment and the RS EEG data acquisition. HP was assessed by the Launay-Slade Hallucination Scale (LSHS), a 16-item questionnaire designed to probe hallucinatory predisposition in the general population (Launay & Slade, 1981). All items were answered on a 5-point Likert scale (0 = “Certainly does not apply to me”, 1 = May not apply to me”, 2 = “Unsure”, 3 = May apply to me”, 4 = “Certainly applies to me”). The total score (sum of all scores on each individual item) ranges between 0-64, with higher scores indicating higher HP. For each participant, we also obtained scores of the 5-item auditory HP (A-HP) subscale and the 3-item auditory verbal HP (AV-HP) subscale following Pinheiro et al. (2020). Fig. 1 shows the distribution of general HP, A-HP, and AV-HP in our sample.

**Fig. 1.**
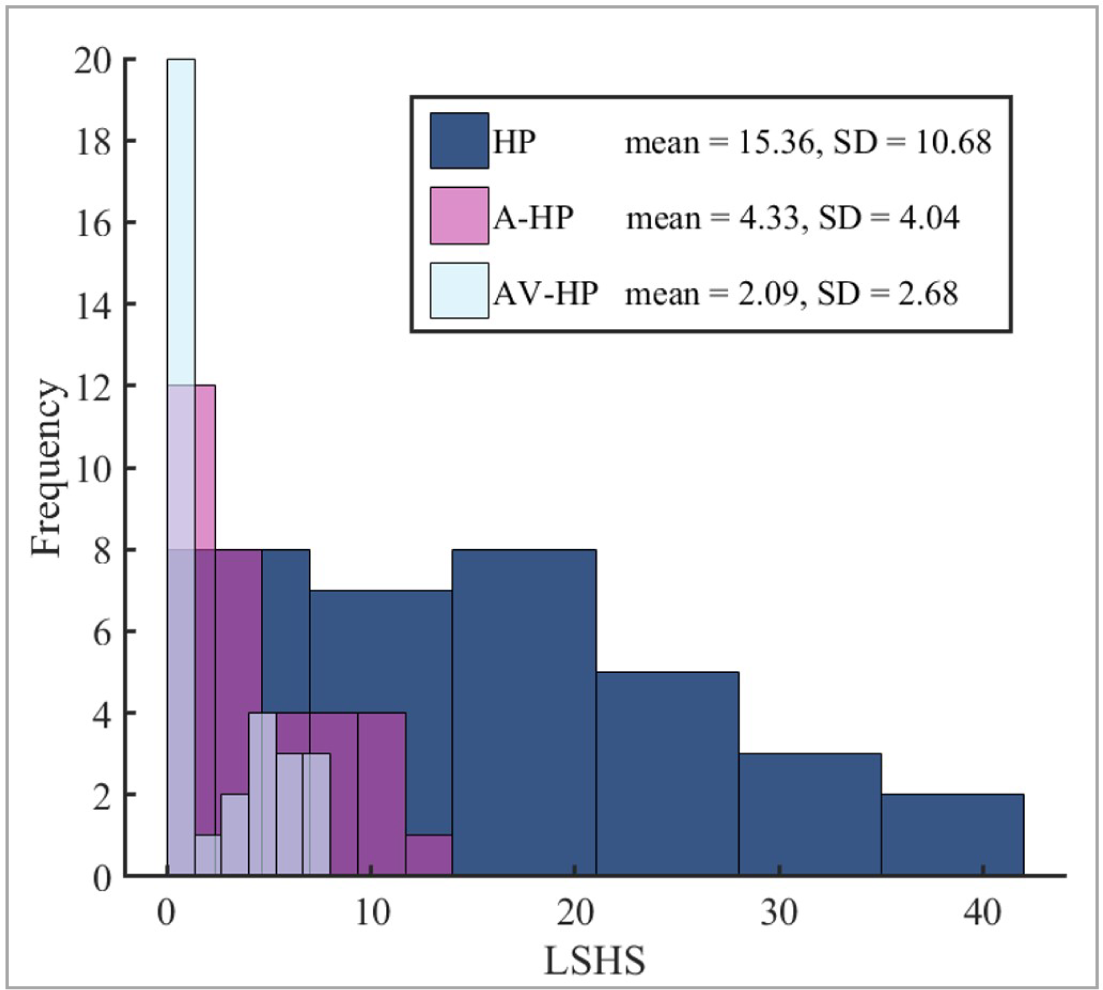
Histograms of total hallucination proneness, auditory hallucination proneness, and auditory verbal hallucination proneness of all participants. HP = Hallucination proneness; A-HP = Auditory HP; AV-HP = Auditory verbal HP.

### EEG recording and processing

Ten minutes (5 minutes eyes-open, EO; 5 minutes eyes-closed, EC; same order for all participants) of RS EEG data were recorded using a 128-channel actiCHAMP active system (Brain Products GmbH, Gilching, Germany) with a sampling rate of 1000 Hz while participants were sitting comfortably in an acoustically shielded EEG-booth. Electrode impedances were kept below 10 kΩ. Participants were asked to stay awake and to minimize body movements including blinking.

EEG data were preprocessed using the MATLAB-based toolbox EEGLAB v2021 (Delorme & Makeig, 2004). The continuous RS EEG data were downsampled to 512 Hz and filtered between 1-40 Hz by means of a finite impulse response (FIR) band-pass filter. Subsequently, the default EEGLAB procedure (clean_rawdata, flat-line criterion = 5; channel criterion = 0.8; line noise criterion = 4) for bad channel detection was used to identify and reject noisy channels. On average, 6.39 (SD = 5.208) channels were rejected per participant. Rejected channels were replaced using spline interpolated neighboring channels. Finally, data were re-referenced to the average of all channels. Automatic Subspace Reconstruction (ASR) with a burst criterion of 20 was applied to remove remaining non-stationary artifacts. Lastly, combined PCA/ICA was applied to the EO and EC data separately to control data rank deficiency and reduce dimensionality. Thus, each individual data set was decomposed into 20 independent components. Components representing i) eyeblinks, ii) horizontal and vertical eye movements, iii) remaining noisy channel activity, iv) muscle-related activity, and v) line noise were rejected. On average, 4.64 (SD = 1.03) components were rejected per dataset. Only EC data were used for subsequent data analyses as attention-induced alpha band modulations are stronger during EC than in EO conditions (Wöstmann et al., 2020).

### Training and test datasets

We assessed if changes in brain state temporal dynamics indicated increased vulnerability to AVH. The employed modeling approach requires a “normative” set of states, which we obtained by splitting the data based on LSHS total scores into “lower” (lower 70% of the LSHS scores, N = 26) and “higher” (upper 30% of the LSHS scores, N = 7) hallucination-prone data sets, serving as training and test sets, respectively. This separation criterion was based on two considerations. First, we aimed to establish a set of states that reflects normative FC signatures of the alpha RS to estimate the states’ temporal dynamics. We expected neither anatomical nor corresponding scalp-topographical differences in FC signatures between lower- and higher-prone individuals so that the same topographical state maps could be used for all participants. Second, the HsMM is tuned to find the best-fitting states based on the statistical properties of the input time series. If individuals on the upper end of the HP continuum would show alterations in state switching behavior (consistent with our expectations), performing the model training on all (N = 33) participants could introduce a bias in the estimation of the states’ dynamics parameters, thereby conflating “normative” and “altered” dynamics. However, we emphasize that splitting the data set was *not* to perform a group comparison but to obtain a realistic and robust model solution. Therefore, individual variability in state dynamics and HP of all participants was considered in the statistical analysis. This allowed accounting for the continuum perspective on hallucinatory predisposition.

### Data preparation and feature extraction

We followed the data processing pipeline as described in Baker et al. (2014). See Fig. 2 for a schematic visualization of our pipeline, including major data processing and analysis steps. The data processing was performed in MATLAB v2020b, using the Brain Dynamics Toolbox and custom scripts (Trujillo-Barreto et al., 2019). To prepare the data of the training set, the following steps were applied for each participant separately. The data were first band-pass filtered to the canonical alpha frequency (8-12 Hz) and linearly detrended. Subsequently, the envelope (i.e., the magnitude of the analytic signal) was extracted using Hilbert transformation and then normalized by the global standard deviation across channels per participant. The latter ensured that any HsMM state would not be driven by differences in amplitude between subjects but rather by the temporal dynamics of the data. The data were then logarithmically transformed. To increase computational efficiency, while preserving most of the information, the training data were temporally concatenated, its dimensionality reduced using 30 principal components, retaining 90.125% of the explained variance, and downsampled to 64 Hz.

**Fig. 2.**
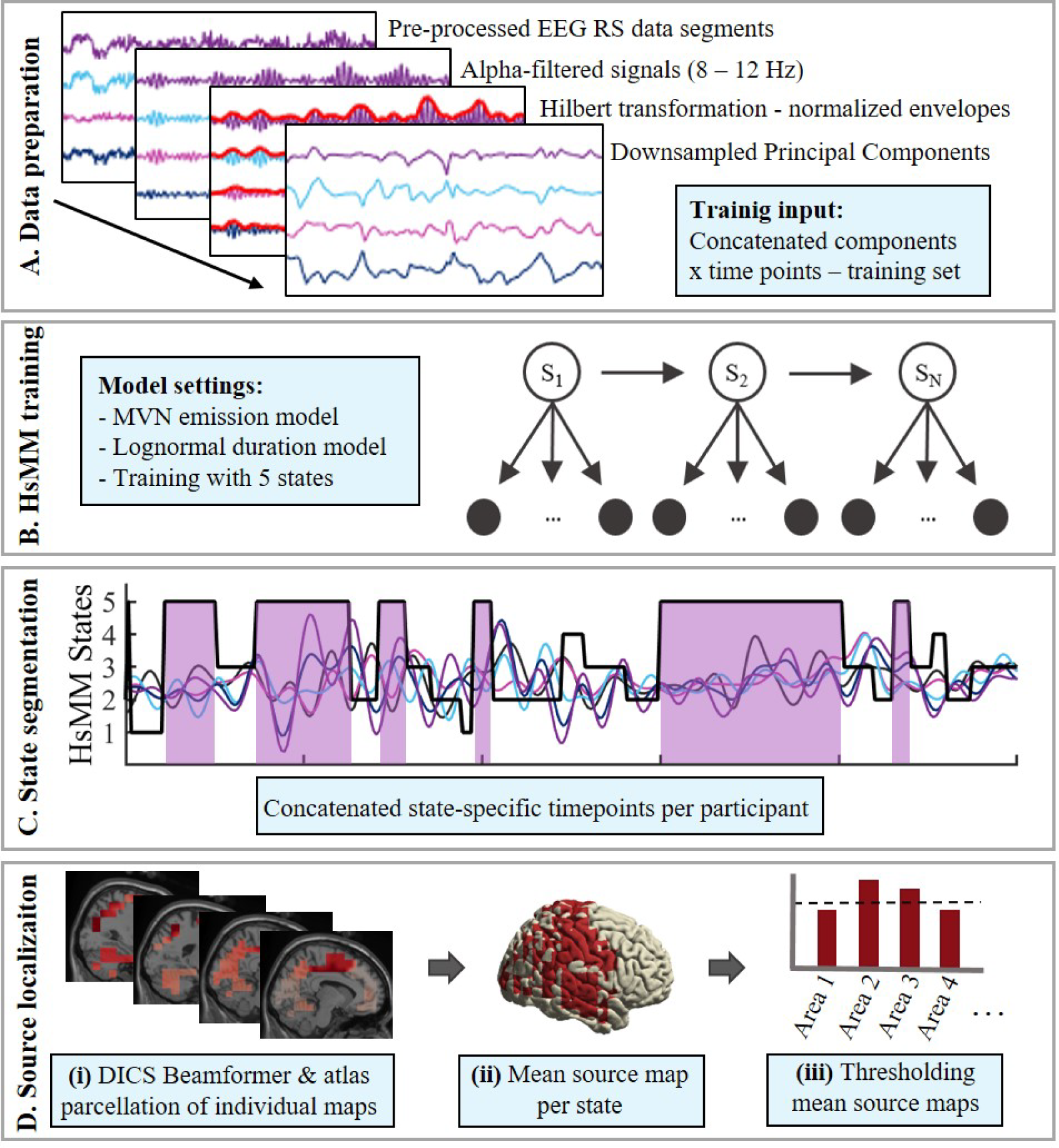
Schematic visualization of the analysis pipeline: Data preparation, HsMM training, and source localization of state-specific data segments. **A. Data preparation.** Pre-processed EEG RS data segments are filtered to the alpha band (8-12 Hz), Hilbert transformed, normalized and log-transformed (log not shown here). Signals were then subjected to PCA with (30 components), downsampled to 64 Hz, and concatenated in time. **B. HsMM training**. Concatenated data were trained using a Multivariate Normal (MVN) emission model and a lognormal duration model to decompose the data into 5 states **C. State segmentation.** Data segments of each state were extracted and concatenated. **D. Source localization**. **i)** Data segments per state and subject were localized using the DICS beamformer implementation in Fieldtrip. Sources were then parcellated into 116 areas using the Automated Anatomical Labeling (AAL) atlas. **ii)** The individual source activity maps were averaged across participants to create mean source maps of each state **iii)** Mean source maps were z-transformed and thresholded to only retain the highest 25% of active sources.

Data of the test set were then transformed accordingly, except that the coefficients obtained from the principal component analysis (PCA) of the training set were used to decompose the data. This prevented data leakage while ensuring that data of both sets were transformed into the same dimensional space to allow valid comparisons of state dynamics across all participants.

### Hidden Semi-Markov Model (HsMM) modeling

Here, we used the Variational Bayes framework of the HsMM described by Trujillo-Barreto et al. (2019). The HsMM is used to model fluctuations in the EEG RS alpha envelope. In this context, a brain state is defined as recurrent and distinct periods of time during which the statistical properties of the data (mean and covariance) of the multichannel envelope are stable (Baker et al., 2014; Trujillo-Barreto et al., 2019). Accordingly, the state transitions mark the points at which the statistical properties of the data change and have been paralleled with the dynamic and systematic switching between underlying RSNs (Baker et al., 2014; Quinn et al., 2018). Importantly, while the states are estimated at the group level (i.e., the states, or recurrent activity patterns are common and fixed for all participants), participant-specific post-hoc state metrics can be obtained directly from the state sequence. This provides insight into individual temporal dynamics of FC, which in turn allow between-participant comparisons.

#### Model inference and initialization

Here, we used a Multivariate Normal (MVN) distribution to model states’ emissions and a log-normal distribution to model the state durations. This implies that the hidden states generate normally distributed data sequences with variable lengths following a log-normal distribution. The variational inference algorithm was randomly initialized and repeated ten times to avoid dependence on initial conditions. Although more repetitions increase the reliability and confidence in the final model, this must be balanced against higher computational costs. The number of states was fixed to five. This number was based on i) preliminary data explorations showing that five states sufficiently characterize between-subject differences in state dynamics, and ii) computational restrictions (i.e., the number of data points that are required to reliably estimate the model parameters). The model was then trained on the data of the 26 lower-scoring participants (i.e., the training set).

#### State sequence, state topographies, and state metrics

The state sequences of all individuals of the training set were directly obtained as the model output. The state sequences of the test set were decoded by applying the already estimated model parameters from the training procedure to the unseen data, which results in a sequence of probabilities of each state being active at each time point. The state with the highest probability was chosen for the state sequence. The states’ topographies were obtained by projecting the mean of the estimated emission distributions from the HsMM back to the sensor space. In other words, the mean vector of the estimated MVN distribution for each state was multiplied by the PCA coefficients obtained during the data processing. This approach is equivalent to the method proposed by Baker et al. (2014). Finally, the mean state duration, i.e., the average time a state is active, and state fractional occupancy (FO), i.e., the total time occupied by a state relative to the total recording time, were extracted to describe the temporal dynamics per state and participant. More specifically, the mean state durations were obtained by fitting a lognormal curve to the histogram of empirical state durations of each subject and state to extract the location (mu) and scale (sigma) parameters of the lognormal distribution. The lognormal mu values were then transformed to a normal scale to obtain interpretable duration values in milliseconds. The FO values were calculated by summing up each individual activation duration divided by the total recording time (i.e., 5 minutes) and are expressed as a percentage.

### Source reconstruction of HsMM states

#### Preparation of state-specific data

To assess whether the HsMM alpha states correspond to well-known RSNs, we used source localization of the original EEG data since the localization of envelope data is not directly interpretable. First, the cleaned RS EEG data were filtered to the alpha band (8-12 Hz) to match the training frequency band. Then, the state sequence (training and test sets; N = 33), concatenated across all participants, was upsampled to match the data sampling frequency after preprocessing/cleaning and prior to data transformation (from 64 Hz to 512 Hz). We chose to upsample the state sequence instead of downsampling the data to keep as many data points as possible to ensure robust source reconstruction. The pre-processed data were then segmented according to the upsampled state sequence into state-specific data points per subject. For each state, the data points of all participants were then standardized and concatenated, resulting in 165 subject-state (33*5) data blocks.

#### Spectral analysis

Spectral analyses and subsequent source localization were performed using the MATLAB-based Fieldtrip toolbox (v20210914) developed at the Donders Institute for Brain, Cognition and Behavior Radboud University, the Netherlands (Baillet, Friston, & Oostenveld, 2011; Oostenveld, Fries, Maris, & Schoffelen, 2011). Multitaper frequency transformation was applied to the full length of each data segment to compute the cross-spectral density (CSD) of the 10 Hz frequency component using a discrete prolate spheroidal sequences (DPSS) taper with a frequency smoothing of 2 Hz.

#### Head model, source model, electrode configuration

We employed a template head (volume conductor) model, normalized to Montreal Neurological Institute (MNI) space, provided by Fieldtrip. The head model was generated using the boundary element method (BEM) (Fuchs, Kastner, Wagner, Hawes, & Ebersole, 2002) and contained meshes for the scalp, skull, and brain with conductivity values of 0.33, 0.0041, and 0.33, respectively. The electrode layout with 128 channels was manually co-registered to the head model using an interactive function in Fieldtrip. The source model was manually generated and contained 2015 vertices spanning the source space with 10 mm resolution. The lead field (i.e., the forward solution), yielding an estimation of how the currents spread from dipolar sources to the sensors, was calculated accounting for the head model and the different tissue conductivities, the pre-defined source space, and aligned electrode configuration.

#### Dynamic Imaging of Coherent Sources (DICS)

Source reconstruction of the subject-state data blocks in the frequency domain was carried out using the Dynamic Imaging of Coherent Sources (DICS) method (Gross et al., 2001) as implemented in Fieldtrip. DICS uses a spatial filter to detect and localize coherent sources, i.e., voxels that show functional synchrony (Drakesmith, El-Deredy, & Welbourne, 2013). In contrast to time-domain beamformer methods, DICS is based on frequency domain data, i.e., the estimation of source coupling relies on the CSD of all sensor pairs. The beamformer output yields a spatial distribution of source power in a chosen frequency band (here alpha). CSD computation and beamformer projection were performed separately for each subject-state data block with regularization of 5%, which is a commonly used regularization strength in the literature and recommended in Fieldtrip. Finally, the beamformer output was normalized by an estimate of the source-projected noise resulting in the Neural Activity Index (NAI) (Van Veen, Van Drongelen, Yuchtman, & Suzuki, 1997).

The individual source images were parcellated using the Automated Anatomical Labeling (AAL) (Tzourio-Mazoyer et al., 2002) into 116 cortical and sub-cortical areas. The parcellated sources were then averaged across participants to obtain one mean source image per state. To assess the areas of highest source activation per state, the mean source images were z-transformed and thresholded to retain only source points exceeding 75-th percentile (z-score = 0.675).

### Statistical analyses

Statistical analyses were performed in IBM Statistics SPSS v26 (IBM Corp., 2019). To assess the predictive value of any of the state’s FO and mean duration values for general HP and the two subscales, A-HP and AV-HP, we performed hierarchical linear regression analyses. We created six regression models: i) HP as dependent variable (DV) and FO values of state 1-5 as predictors, ii) HP as DV and mean duration values of state 1-5 as predictors, iii) A-HP as DV and FO values of state 1-5 as predictors, iv) A-HP as DV and mean duration values of state 1-5 as predictors, v) AV-HP as DV and FO values of state 1-5 as predictors, and finally vi) AV-HP as DV and mean duration values of state 1-5 as predictors. For all models, we used stepwise regression with backward elimination where the criterion for exclusion of predictors was set to the probability of F >/= 0.1.

## Results

### Results of the HsMM inference

Fig. 3 shows FO and mean durations for each participant in panels A and B, respectively. All states were represented in all participants, but their occupancy and mean duration show considerable variability between participants. This confirmed that the states identified by the model were common to all participants and that the HsMM captured individual differences in the state dynamics. Fig. 4 depicts the state maps, i.e., the most representative topography for each state based on the alpha amplitude envelope, along with histograms of FO and mean duration per state. The order of states is arbitrary. State 4 and 5 showed a similar activity distribution, although with opposite polarity, which could represent complementary states of high and low alpha power. However, the FO and mean duration histograms showed quite different dynamics, with state 4 showing longer durations up to 500 ms, whereas state 5 durations are clustered around 200 ms. The mean state durations of all states vary between .17 and .22 seconds.

**Fig. 3.**
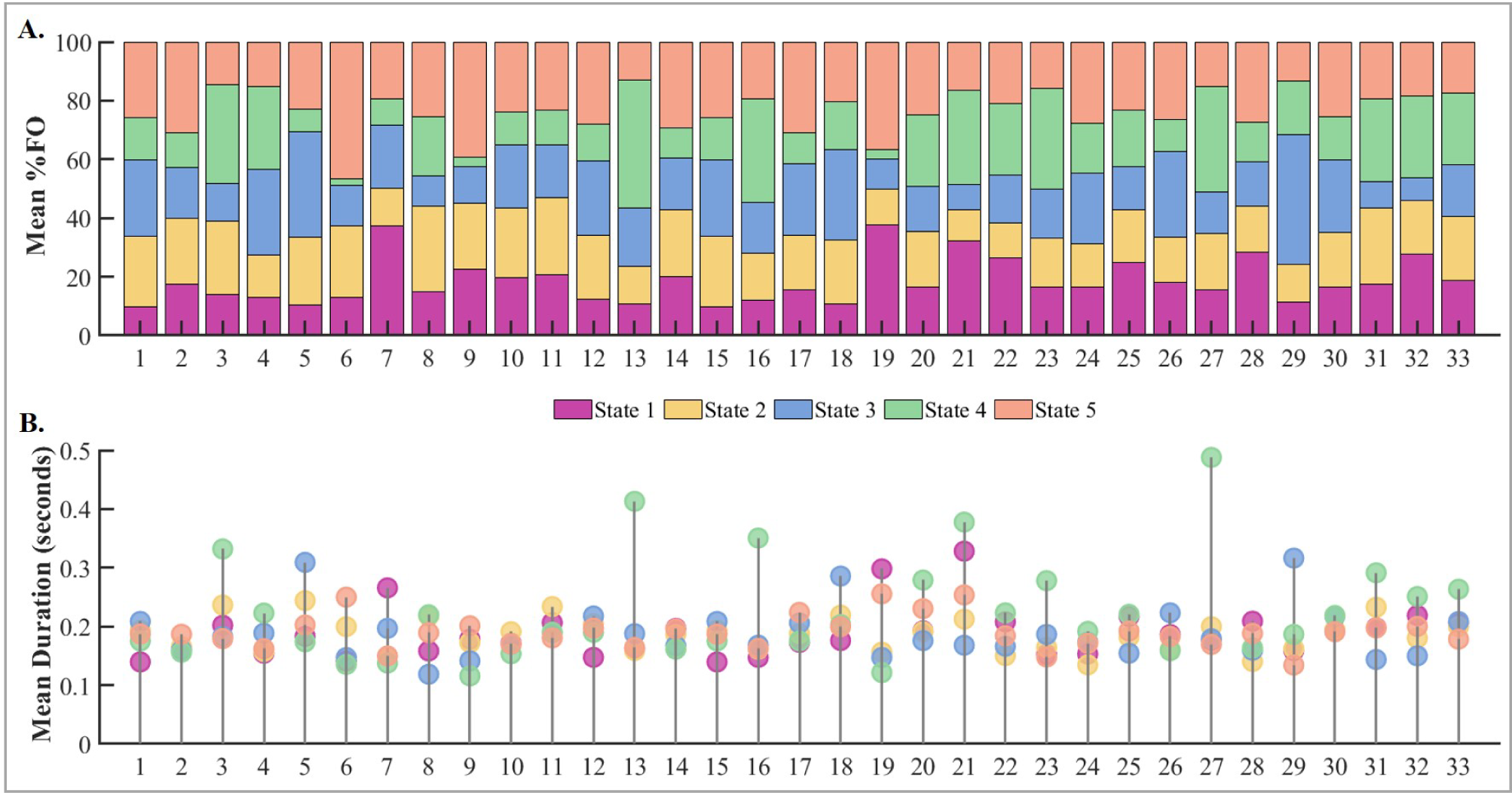
Fractional occupancy and mean duration of all states and participants. Distribution of Fractional Occupancy (FO, panel A) and mean duration (panel B) of ‘lower’ hallucination-prone (HP) participants (1-26) and decoded ‘higher’ HP participants (27 – 33). FO is provided in percentage (%). Durations are given in seconds.

**Fig. 4.**
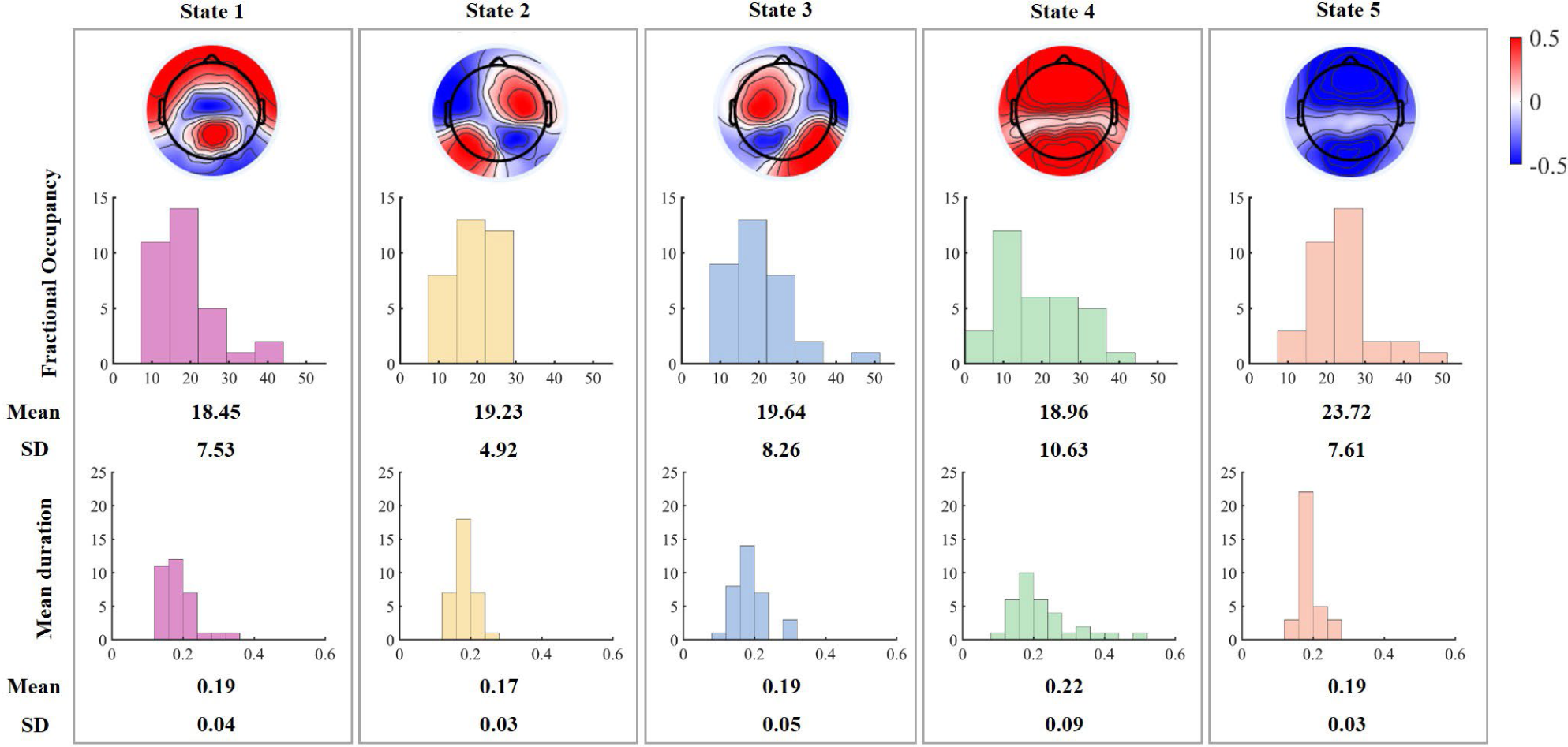
The maps depict the most representative topography for each state based on the alpha amplitude envelope. The order of the states is arbitrary. The scale of the state maps represents microvolts. The upper and lower panels show histograms of FO and mean duration per state, respectively.

### Results of the statistical analyses

Assumptions such as normality of residuals, linearity, and homoscedasticity for the regression models for HP and A-HP as DVs were met. For the models with AV-HP as DV, the residuals were not normally distributed, which may compromise the generalizability of the results. The results of the final significant models (i.e., after the exclusion of insignificant predictors) are summarized in Table 1. The complete results of all regression models including all steps of the backward elimination procedure can be found in the supplementary material (section B, supplementary tables 2-7). The stepwise regression with A-HP as DV resulted in a model with only state 1 FO as significant predictor F(1,32) = 7.818, p = .009, which accounted for 20.1 % of the variation in A-HP (R^2^ = .201). Similarly, the model with A-HP as DV and duration values as predictors indicated state 1 mean duration as only significant predictor F(1,32) = 5.949, p = .021, accounting for 16.1% of the variation (R^2^ = .161). Further, the final regression model with AV-HP as DV also revealed state 1 FO as significant predictor, F(1,32) = 5.219 p = .029, which accounted for 14.4 % of the variation in AV-HP (R^2^ = .144). Finally, the mean duration of state 1 significantly predicted AH-HP, F(1,32) = 5.070 p = .032, explaining 14.1 % of the variation in AH-HP (R^2^ = .141). The two regression models with HP as DV did not reveal any significant predictors.

**Table 1.**
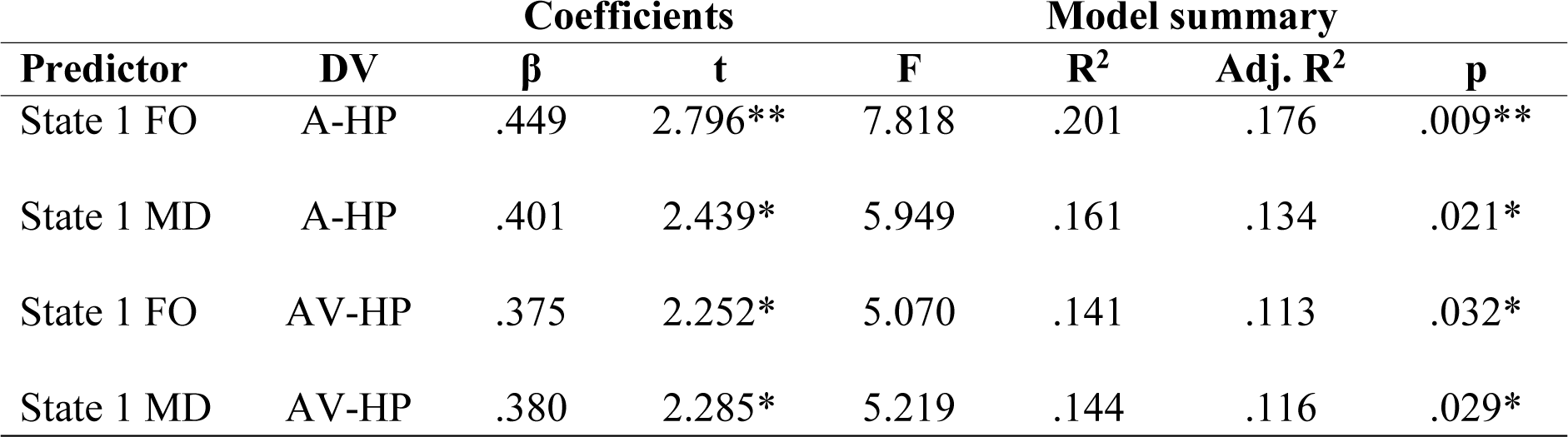
Significant model results of the regression analyses; N = 33; *p < .05, ** p < .01. DV = dependent variable, FO = fractional occupancy, MD = mean duration. A-HP = Auditory hallucination proneness, AV-HP = Auditory verbal hallucination proneness. Each row represents the final model for each stepwise regression analysis.

In summary, state 1 FO and mean duration significantly predicted both A-HP and AV-HP, but not general HP. That means that individuals scoring higher on the auditory and auditory-verbal subscales of the LSHS showed greater occupancy and longer duration of state 1. This might indicate an effect specific to the auditory modality of a hallucinatory experience. The explained variance of the final models ranged between 14.1 - 20.1%.

### Results of the source localization

Fig. 5 shows the mean source images, i.e., the cortical distribution of active sources per state. All states showed a similar source activity distribution, localized in the posterior part of the brain, spanning the superior and inferior parietal cortex, posterior cingulate cortex (PCC), the cuneus and precuneus, parts of the operculum and of the occipital cortex. However, despite overall similarity, there were some noticeable differences. State 1 source activity extended to bilateral superior and middle temporal cortices, whereas state 2 and 3 were more left-lateralized. State 4 was characterized by source activity in the posterior parietal cortex bilaterally and in the right inferior temporal lobe. State 5 again showed a more asymmetrical activation pattern and was defined by source activity in the left, but not the right, inferior and superior parietal cortex. Additionally, state 5 showed more activation in the right temporal lobes, as compared to the left. The labels of all active areas surviving the thresholding per state are listed in the Supplementary Material (Section A - Supplementary Table 1). For an orthogonal and slice view of the mean activation per state, please refer to Supplementary Figures 1 and 2, respectively).

**Fig. 5.**
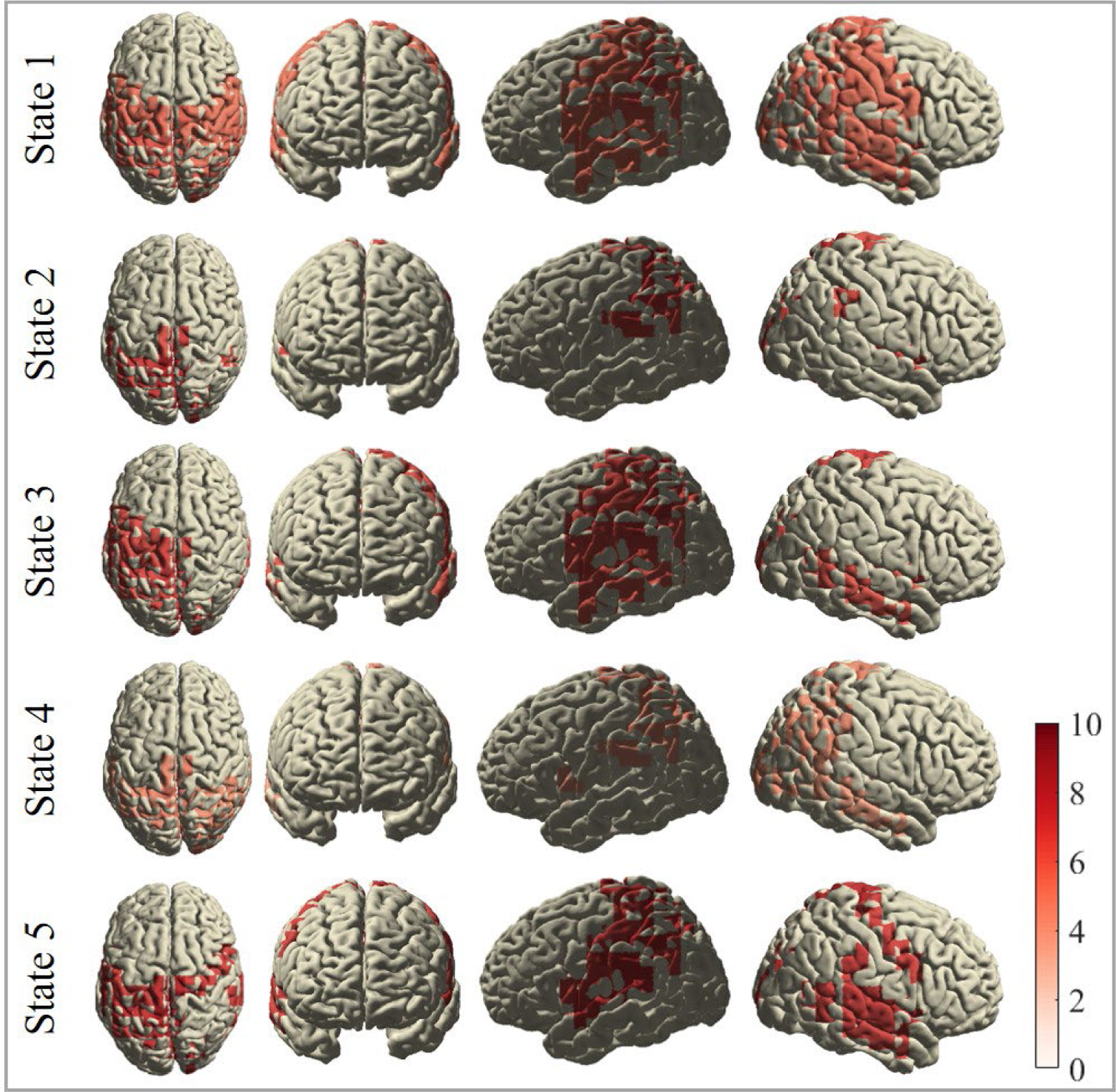
Cortical distribution of active sources (parcellated mean source images per state). Activation is expressed as Neural Activity Index (NAI) (Van Veen et al., 1997). Images were generated by i) parcellating the individual source images using the AAL atlas into 116 areas, ii) averaging the activity within each parcel across participants, iii) thresholding the mean sources images based on the 75-th percentile of source activity (>/= z-value of 0.675), and iv) interpolating the active sources onto a template brain mesh.

## Discussion

This study investigated if brain state dynamics of alpha band activity recorded at rest predict non-clinical HP, A-HP, and AV-HP. To this end, five brain states and their dynamics features were estimated by means of an HsMM. Using frequency-domain source reconstruction, we further assessed the source activity of each state and whether they overlapped with well-known RSNs. used to predict HP, A-HP, and AV-HP using stepwise linear regression. The results showed that the HsMM successfully captured the temporal complexity of narrow-band alpha RS activity and characterized brain states with distinct temporal and spatial profiles. Moreover, mean duration and FO of a state corresponding to auditory, sensorimotor, and DMN areas (here state 1) predicted individual differences in A-HP and AV-HP, but not general HP. These findings confirm that the HsMM approach is sensitive to subtle brain state dynamics and can reveal systematic physiological spatio-temporal patterns that link to cognitive and behavioral states (Honcamp et al., 2022; Trujillo-Barreto et al., 2019). The results of the source localization further corroborate previous findings that linked alpha band activity to posterior hubs of the DMN (Hindriks et al., 2017; Mantini, Perrucci, Del Gratta, Romani, & Corbetta, 2007). Despite the states’ overall similarity in the source domain, the states showed some spatial specificity in temporal and subcortical brain structures, confirming temporally and spatially distinct patterns of underlying network FC. Given that the identified HsMM states reflect recurrent signatures in alpha band activity independent of task demands (Craddock et al., 2017; Klimesch et al., 2007; Wöstmann et al., 2020), the state’s temporal dynamics could inform about spontaneous shifts in attention, awareness, and perceptual sensitivity, rather than perceptual accuracy per se. Accordingly, state 1 may reflect an attentional bias toward internally generated events and heightened awareness of and sensitivity for auditory sensations. Together, this could suggest increased vulnerability to hallucinatory experiences in the auditory domain.

### Resting state alpha-band dynamics and perceptual sensitivity

Defining the role of alpha band activity in the resting EEG is not straightforward. This is partly due to the presence of multiple rhythms in the alpha range, including the visual alpha rhythm, the somatosensory mu-rhythm, as well as the temporal tau and breach rhythms. As these rhythms stem from distinct (sub-)cortical generators, they are hypothesized to serve different cognitive functions (Manshanden, De Munck, Simon, & da Silva, 2002; Niedermeyer, 1990; Tiihonen et al., 1991). Despite this differentiation, a consistent finding is the particularly strong RS alpha power in posterior and post-central cortices including occipital, somatosensory, and temporal regions (Hindriks et al., 2017). The presence of several alpha rhythms may also explain that the identified HsMM states are characterized by different combinations of active sources. This emphasizes the spatial complexity of the alpha band activity during RS (Ben-Simon, Podlipsky, Arieli, Zhdanov, & Hendler, 2008; Hindriks et al., 2017).

In cognitive tasks, alpha band activity is linked to selective attention, cognitive control, and perceptual sensitivity (Hindriks et al., 2017; Klimesch et al., 2007). Other studies found that pre-stimulus alpha power is inversely related to perceptual awareness but does not predict discrimination accuracy (Benwell, Coldea, Harvey, & Thut, 2022; Benwell et al., 2017). Similarly, Iemi and Busch (2018) suggested that alpha power fluctuations link to changes in neural baseline excitability and thus influence neural responsiveness to both signal and noise. In turn, alpha band activity dynamics may reflect criterion changes in an individual’s response behavior (i.e., heightened neural excitability leading to a more liberal responses), rather than perceptual accuracy (Iemi & Busch, 2018). Consistent with these findings, parieto-occipital alpha band activity is associated with perceptual stability when exposed to ambiguous sensations (Helfrich et al., 2016; Katyal, He, He, & Engel, 2019). These results support the role of alpha band activity in the processing of sensory inconsistencies. However, most of these findings were task-related and are not necessarily transferable to spontaneous alpha band fluctuations during RS. Interestingly, during pre-task RS, alpha fluctuations predict situational awareness in a subsequent task (Kaur, Chaujar, & Chinnadurai, 2020). This suggests that task-independent alpha oscillations can shape information processing by selectively modulating awareness and attention to internal and external stimuli. Although a direct effect of alpha modulations on cognition is difficult to show in pure RS conditions, evidence supports the role of alpha in alertness, attention, cognitive control, and perceptual stability (Braboszcz & Delorme, 2011; Katyal et al., 2019; Mahjoory, Cesnaite, Hohlefeld, Villringer, & Nikulin, 2019; Mathewson, Gratton, Fabiani, Beck, & Ro, 2009; Sadaghiani & Kleinschmidt, 2016).

In the RS, the brain engages in non-sensory activities like mind-wandering and autobiographical memory retrieval (Raichle, 2015). It was shown that spontaneous cognitive operations during RS can be broken down into different cognitive modes (referred to as phenotypes, e.g., theory of mind, planning, sleepiness) with inter-individual differences in shifting patterns and individual preferences for a given mode (Diaz et al., 2013). These RS phenotypes are characterized by distinct electrophysiological signatures of cognition and behavior (Pipinis et al., 2017; Tarailis, Šimkutė, Koenig, & Griškova-Bulanova, 2021). For example, the Somatic Awareness RS phenotype is associated with the class C microstate, which in turn, has been linked to psychosis (Pipinis et al., 2017). Interestingly, microstates C and D were suggested as endophenotypes of schizophrenia, as patients as well as their siblings, showed similar microstate temporal dynamics (da Cruz et al., 2020). However, directly comparing microstates and HsMM states is challenging. While the state topographies derived by the microstate analysis and the HsMM could be comparable, the state switching patterns are quite distinct and suggest that the temporal dynamics of the time series are differently encoded in the two approaches (Rukat et al., 2016). Similarly, Coquelet et al. (2021) showed that, unlike microstates, the HMM (and likely also HsMM) states are not only driven by power fluctuations but also by the cross-covariance of the multivariate emissions, i.e., FC fluctuations over time. Thus, the H(s)MM accounts for critical temporal features that the microstates dismiss.

We conclude that the HsMM state sequence could reflect shifts between different cognitive modes during rest that might coincide with modulations in selective attention between specific sensory modalities and sensory responsiveness. Accordingly, the transitions between attentional states might correspond to sequential changes in brain activation patterns. This could explain the observed spatial differences in active sources and source power between HsMM states. Thus, the identified alpha-band HsMM states could be understood as unique combinations of active sources that correspond to the same (overlapping) alpha generators resulting in distinct FC states. The states could further reflect inter-individual differences in attentional switches between sensory modalities that are characterized by distinct temporal dynamics such as FO and state duration.

Interestingly, the sources of our alpha-band HsMM states show similarities with the “posterior higher-order cognitive” state identified by Quinn et al. (2018), characterized by posterior nodes of the DMN and high coherence and power in the alpha frequency. It must be noted that their approach considerably differs from ours, in that Vidaurre et al. (2018) used time-domain source-reconstructed MEG data as input to a time-delay embedded HMM (TDE-HMM). The TDE-HMM allows identification of both spectrally and temporally resolved information about the identified states. However, both approaches confirm that spontaneous brain activity is organized into short-lived recurrent brain states with specific spectral information and spatial correspondence to well-known RSNs.

### Relevance for hallucination proneness

Recent studies show that hallucinatory experiences with varying severity in the general population may be much more common than previously reported (Linszen et al., 2022). This emphasizes the need for a better understanding of their neural correlates. The current results suggest that RS alpha band dynamics might reflect individual differences in hallucinatory predisposition. Alpha band activity modulations, especially in eyes-closed conditions, were suggested to reflect changes in auditory attention through the inhibition of irrelevant, non-salient sensory information (Strauß, Wöstmann, & Obleser, 2014; Wöstmann et al., 2020). These findings support the hypothesis that alpha band dynamics shape the allocation of attentional resources (Jensen, Bonnefond, & VanRullen, 2012). Alternatively, the relationship between HP and alpha band dynamics may be explained by modulations of neural excitability (Iemi & Busch, 2018). Heightened baseline excitability is thought to amplify both signal and noise, which may alter an individual’s tendency to “detect” stimuli in the absence of sensory stimulation and confuse internally generated sensory events as coming from an external source (Iemi & Busch, 2018; Ilankovic et al., 2011; Northoff & Qin, 2011; Stephane, Kuskowski, McClannahan, Surerus, & Nelson, 2010). Interestingly, the relationship with state 1 occupancy and mean durations was only found for the auditory and auditory-verbal modality of hallucinatory experiences but not for general HP. This raises the question of whether alpha band dynamics not only reflect fluctuations in attention and neural responsiveness to internally generated information, but also how attentional resources are distributed between sensory modalities as suggested by Keller, Payne, and Sekuler (2017).

The source localization of state 1 revealed simultaneously active sources located in bilateral pre-, para-, and post-central lobes, bilateral inferior and superior parietal lobes, bilateral (pre-)cuneus, bilateral PCC, and bilateral superior and middle temporal lobes. These regions are associated with the somatosensory network, posterior hubs of the DMN, and the auditory network (Damoiseaux et al., 2006; Raichle, 2015). This finding suggests that brain states, as estimated from H(s)MMs, associate with a mixture of RSNs and represent activity configurations driving cognitive processes (Chen, Langley, Chen, & Hu, 2016; Hunyadi et al., 2019). Michael, Salgues, Plancher, and Duran (2022) found that spontaneous oscillatory activity in sensorimotor cortices reflects individual differences in bodily awareness. One may speculate that such activity not only indicates distortions of bodily perception but potentially also delusions and hallucinations. The DMN has been associated with spontaneous, task-independent cognition, including mind-wandering and daydreaming (Raichle, 2015). The PCC, a posterior hub of the DMN, is thought to play a major role in internally directed cognition such as autobiographical memory, but also in directing and reorienting attention. Further, functional interactions between the PCC and other brain networks are crucial for conscious awareness and perception (Leech & Sharp, 2014). Activation of the superior and middle temporal lobes is commonly associated with the processing of incoming auditory stimuli. Abnormal engagement of both the primary and secondary auditory cortices in the absence of sensory stimulation has been related to the experience of AVH (Kompus, Westerhausen, & Hugdahl, 2011; Northoff & Qin, 2011). It is important to note that such findings often come from so-called symptom capture studies, which measure AVH-related brain activity while asking participants to indicate the on- and offset of hallucinations by a button press. Thus, the recorded brain activity might be confounded by directed attention to the hallucinations and not reflect the brain’s true RS. Along with aberrant auditory cortex activation, many studies have reported abnormally elevated DMN activation in individuals with AVH, which may indicate an attentional bias toward internally generated (auditory) events and a failure in downregulating auditory processing systems (Kompus et al., 2011; Northoff & Qin, 2011). Schizophrenia patients with AVH further exhibit increased connectivity between posterior parts of the DMN and regions associated with auditory processing, which may contribute to the misattribution of self-generated sensory events to an external source, and thus the experience of auditory hallucinations (Mannell et al., 2010; Northoff & Qin, 2011). Similarly, a fMRI network analysis revealed that aberrant activation of the DMN and the auditory network links to the predisposition to hallucinate in non-psychotic individuals (van Lutterveld et al., 2014), suggesting that changes in DMN dynamics are not unique to the clinical population but may serve as an early marker of increased HP in the general population. Lastly, Kottaram et al. (2019) applied an HMM to fMRI BOLD hemodynamics to investigate differences in RSN dynamics between patients with schizophrenia and healthy control participants. They found that patients spent a significantly shorter proportion of time in a state characterized by high DMN and low sensorimotor network activation, however, once visited, the duration of that state was significantly longer as compared to the controls. The authors concluded that schizophrenia is characterized by a reduced dynamism of the DMN, especially regarding its interaction with sensory networks. The disturbed DMN dynamics further correlated with the severity of positive symptoms, including hallucinations.

The participants tested in the current study were non-hallucinating individuals varying in HP. According to the continuum perspective, high hallucination-prone individuals without AVH should show attenuated changes in brain activity as compared to non-clinical and clinical voice hearers (Johns et al., 2014; Johns & Van Os, 2001; van Lutterveld et al., 2014). In summary, the current results show that non-clinical A-HP and AV-HP link to changes in alpha-band temporal dynamics of a state that is characterized by posterior DMN and auditory network activation. Thus, individuals who are prone to auditory (verbal) hallucinatory experiences spend longer time segments in a state that may reflect an increased bias toward internal events, potentially as they are more salient by default (Kapur, 2003), combined with heightened neural responsiveness to (internally generated) auditory percepts.

### Limitations and Future Directions

To investigate temporal dynamics as potential neural correlates of the non-clinical HP continuum, the study sample comprised participants without AVH but with varying levels of HP. Although the continuum perspective of psychosis-like symptoms has gained considerable evidence through different methodological approaches, it does not fully explain the heterogeneity of neurocognitive changes and phenomenological characteristics in non-clinical and clinical AVH (Badcock & Hugdahl, 2012; Corona-Hernández, Brederoo, de Boer, & Sommer, 2022). In comparison to non-clinical AVH, clinical voice hearers commonly report their AVH to be negative, associated with low controllability, and accompanied by other symptoms such as delusions (de Leede-Smith & Barkus, 2013; Larøi et al., 2012; Sommer et al., 2010). Thus, it is questionable whether such qualitative differences can be explained by the same continuum of neural changes or whether so-called quasi-dimensional models of psychosis better account for this heterogeneity (Baumeister, Sedgwick, Howes, & Peters, 2017; Corona-Hernández et al., 2022). For instance, analysis of the functional connectome suggests that different neural mechanisms may underlie hallucinatory experiences across the psychosis continuum in non-clinical and different clinical populations, e.g., patients with schizophrenia and bipolar disorder. (Schutte et al., 2022). Future research should therefore explore whether similar changes in RS temporal dynamics are also characteristic of individuals with AVH in the non-clinical and clinical spectrum. Additionally, future studies could apply the introduced approach to hallucinations in another, e.g., the visual domain, to explore modality-specific state changes. Extending the current approach to the whole HP continuum as well as multiple sensory modalities could help advance theories about psychosis proneness, pre-clinical risk analysis, and interventions.

Of further note is that the current approach aimed at deriving a normative set of states, common to all participants, and considering inter-individual differences in their temporal dynamics. Splitting data into a training set and a test set is a common modeling approach to model the extracted features of several data sets and subsequently to evaluate out-of-sample generalization performance of the modeling procedure (Murphy, 2012). Such methods have been applied to RS EEG data in various clinical contexts (Chen, Lu, Xie, & Shang, 2020; de Miras, Ibáñez-Molina, Soriano, & Iglesias-Parro, 2023; Huggins et al., 2021; Mao, Zhu, Li, Zhang, & Sun, 2018; Xu, Zheng, Mao, Wang, & Zheng, 2020). Moreover, a similar splitting of the LSHS scores into higher and lower scoring individuals was used in prior studies (Collignon, Van der Linden, & Larøi, 2005; Garrison et al., 2017; Kanemoto, Asai, Sugimori, & Tanno, 2013; Larøi, Van der Linden, & Marczewski, 2004). While the current approach is justified for its specific aim, it might be worthwhile to explore if and how the model solution and the state’s temporal dynamics change as a function of the splitting criterion. Such an exploration could inform about current perspectives of the HP continuum, e.g., whether (i) the same set of states can be used to describe underlying neurophysiological phenomena for the whole HP continuum or (ii) differences between non-clinical and clinical voice hearers are better accounted for by a distinct (or overlapping) set of states.

The modeling of temporal dynamics in the current study was restricted to the alpha band, in accordance with its role in selective attention, cognitive control, and perceptual sensitivity, as well as its association with DMN activation (Craddock et al., 2017; Hillebrand et al., 2012; Hindriks et al., 2017). This narrow-band filtering approach may also have influenced the spatial overlap between states on the level of the sources. However, it is important to consider that the identified active sources, e.g., the PCC, do not function in isolation but dynamically interact with anatomically and functionally connected brain areas in more distant parts of the brain, e.g., the medial prefrontal cortex, another major hub of the DMN. Additionally, broadband data may reflect a richer repertoire of the electrophysiological signatures of the underlying RSNs operating within different frequency bands (Mantini et al., 2007). Thus, our current approach yields a specific view on RS dynamics through the lens of the alpha band frequency. Future studies could therefore explore how brain state temporal dynamics enfold in broad-band data and how they differ between frequency bands. This could offer further insights into the predictive value of frequency-specific RS alterations for cognition and behavior. As the current results emphasize the non-stationarity of the RS, additional quantification of RS phenotypes as suggested by Diaz et al. (2013) could facilitate the interpretation of brain states and inform about cognitive biases.

Lastly, it is important to note that the current focus was on temporal dynamics and key sources of alpha brain states as a function of HP. Accordingly, the chosen approach revealed simultaneously and highly active sources of each state but did not provide evidence for functionally connected large-scale brain networks. Future research should therefore aim to unveil interaction between (sub-)network dynamics along the whole HP continuum. Adaptations of the H(s)MM method using a multivariate autoregressive (MAR) emission model (Hernandez et al., 2022; Vidaurre et al., 2016) can be used to model effective connectivity based on both amplitude and phase dynamics. Such models could yield important further insights into RSN interactions and their relationship with HP non-clinical and clinical populations.

## Conclusions

The brain’s RS fluctuates between distinct FC states on a sub-second timescale that reflects the interactions of large-scale brain networks. In the current study, we used a HsMM to characterize temporal dynamics of alpha band states and showed that increased occupancy and mean duration of state 1 are associated with non-clinical auditory and auditory-verbal HP, but not general HP. This suggests that changes in RS dynamics link to altered cognition and behavior. The localization of state 1 in posterior parts of the DMN and parts of sensorimotor and auditory networks is consistent with earlier research on RSN-related changes in clinical and non-clinical voice hearers (Northoff & Qin, 2011; Van Diessen et al., 2015). These findings suggest that increased attentional bias towards internally generated auditory events, as it is often found in clinical voice-hearers, might already be characteristic of healthy but high-hallucination-prone individuals. In line with previous studies, these results underline the role of alpha band dynamics in perceptual sensitivity and the processing of ambiguous sensory information (Helfrich et al., 2016; Iemi & Busch, 2018; Katyal et al., 2019). We conclude that the HsMM is a sensitive new approach that can characterize narrow-band RS fluctuations on a sub-second timescale and help derive meaningful neurophysiological correlates of the HP continuum.

## Author contributions

HH: Conceptualization, recruitment and data collection, development of the data analysis pipeline, data analysis, manuscript writing, editing

SXD: Recruitment and data collection, editing

JRC: Development of the data analysis pipeline, data analysis, editing

AA: Development of the data analysis pipeline; editing

WeD: Conceptualization, development of the data analysis pipeline, editing

NTB: Software implementation, conceptualization, development of the data analysis pipeline, editing

MS: Conceptualization, development of the data analysis pipeline, manuscript writing, editing

SK: Conceptualization, development of the data analysis pipeline, manuscript writing, editing

DL: Conceptualization, editing

TvA: Conceptualization, editing

## Funding information

HH is funded by the Studienstiftung des Deutschen Volkes, Germany. SK and MS are funded by the BIAL foundation [BIAL 146/29].

WeD acknowledges ANID, Chile [projects FONDECYT 1201822, ANILLO ACT210053 and Basal FB0008], ValgrAI and the Generalitat Valenciana, Spain.

NTB acknowledges the Engineering and Physical Sciences Research Council (EPSRC) [EP/N006771/1] and the Medical Research Council (MRC) [MR/X005267/1], UK.

## Data availability statement

The software used for HsMM implementation in this manuscript is available at ttps://github.com/daraya78/BSD. Anonymized data will be shared with other researchers upon reasonable request to the corresponding author.

## Acknowledgements

The authors would like to thank Alexandra Kalberer and Maren Cremer for their assistance in data collection.

## Ethics approval statement

Ethical approval was granted by dutch Medisch-ethische toestingscommissie (METC 20-035).

## Participant consent statement

Participants provided written informed consent prior to study participation.

