## Supplementary Material for "EEG resting state alpha dynamics predict individual proneness to auditory hallucinations"

#### A. Active sources per state

**Supplementary Table 1.** Active sources after thresholding (highest 25 %) per state

| State 1 | State 2 | State 3 | State 4 | State 5 |
|---|---|---|---|---|
| Frontal Inf Oper L | Rolandic Oper R | Precentral L | Frontal Inf Oper L | Precentral R |
| Rolandic Oper L | Cingulum Mid L | Frontal Inf Oper L | Rolandic Oper L | Frontal Inf Oper L |
| Rolandic Oper R | Cingulum Mid R | Rolandic Oper L | Cingulum Mid L | Rolandic Oper L |
| Cingulum Mid L | Cingulum Post L | Rolandic Oper R | Cingulum Mid R | Rolandic Oper R |
| Cingulum Post L | Cingulum Post R | Cingulum Mid L | Cingulum Post L | Cingulum Mid L |
| Cingulum Post R | Hippocampus R | Cingulum Post L | Cingulum Post R | Cingulum Post L |
| Calcarine R | Calcarine R | Cingulum Post R | Calcarine R | Cingulum Post R |
| Cuneus L | Cuneus L | Calcarine L | Cuneus L | Calcarine R |
| Cuneus R | Cuneus R | Calcarine R | Cuneus R | Cuneus L |
| Occipital Sup R | Lingual R | Cuneus L | Lingual L | Cuneus R |
| Occipital Mid R | Occipital Sup R | Cuneus R | Occipital Sup R | Occipital Sup R |
| Postcentral L | Fusiform R | Lingual L | Occipital Mid R | Postcentral L |
| Postcentral R | Parietal Sup L | Lingual R | Parietal Sup L | Parietal Sup L |
| Parietal Sup L | Parietal Inf L | Occipital Sup R | Parietal Sup R | Parietal Inf L |
| Parietal Sup R | Parietal Inf R | Postcentral L | Parietal Inf L | SupraMarginal L |
| Parietal Inf L | SupraMarginal L | Parietal Sup L | Parietal Inf R | SupraMarginal R |
| Parietal Inf R | Angular L | Parietal Inf L | SupraMarginal L | Angular L |
| SupraMarginal L | Precuneus L | SupraMarginal L | Angular L | Precuneus L |
| SupraMarginal R | Precuneus R | Angular L | Angular R | Precuneus R |
| Angular L | Paracentral Lobule L | Precuneus L | Precuneus L | Paracentral Lobule L |
| Angular R | Paracentral Lobule R | Precuneus R | Precuneus R | Paracentral Lobule R |
| Precuneus L | Thalamus L | Paracentral Lobule L | Paracentral Lobule L | Thalamus L |
| Precuneus R | Thalamus R | Paracentral Lobule R | Paracentral Lobule R | Heschl L |
| Paracentral Lobule L | Heschl R | Caudate L | Caudate L | Heschl R |
| Paracentral Lobule R | Cerebellum 4 5 L | Pallidum L | Putamen L | Temporal Sup L |
| Caudate L | | Thalamus L | Pallidum L | Temporal Sup R |
| Thalamus L | | Heschl L | Thalamus L | Temporal Mid R |
| Heschl L | | Heschl R | Thalamus R | |
| Heschl R | | Temporal Sup L | Heschl R | |
| Temporal Sup L | | Temporal Mid L | Temporal Mid R | |
| Temporal Sup R | | Temporal Mid R | | |
| Temporal Mid L | | Vermis 1 2 | | |
| Temporal Mid R | | Vermis 3 | | |

#### B. Complete results of the hierarchical linear regression models (including non-significant models)

This section contains the complete results of the regression analyses. For all models, we used a backward exclusion procedure in which non-significant predictors were removed stepwise according to the following criterion: Probability of F-to-remove >/= 0.1.

**Supplementary Table 2.** General HP as dependent variable and FO values of states 1-5 as predictors.

| | Coefficients | | | Model summary | | | |
|---|---|---|---|---|---|---|---|
| **Predictor** | **β** | **t** | **p** | **F** | **$R^2$** | **Adj. $R^2$** | **$R^2\Delta$** |
| **Step 1** | | | | .581 | .077 | -.055 | .077 |
| (Constant) | - | 1.821 | .079 | | | | |
| State 1 FO | -.077 | -.283 | .779 | | | | |
| State 2 FO | -.281 | -1.151 | .259 | | | | |
| State 3 FO | -.189 | -.796 | .433 | | | | |
| State 5 FO | -.25 | -.128 | .899 | | | | |
| **Step 2** | | | | .797 | .076 | .11 | -.001 |
| (Constant) | - | 1.849 | .075 | | | | |
| State 1 FO | -.086 | -.330 | .744 | | | | |
| State 2 FO | -.292 | -1.30 | .204 | | | | |
| State 3 FO | -.191 | -.814 | .420 | | | | |
| **Step 3** | | | | 1.175 | .073 | .073 | -.003 |
| (Constant) | - | 3.148 | .004 | | | | |
| State 2 FO | -.248 | -1.398 | .172 | | | | |
| State 3 FO | -.142 | -.800 | .430 | | | | |
| **Step 4** | | | | .230 | .053 | .022 | -.020 |
| (Constant) | - | 3.319 | .002 | | | | |
| State 2 FO | -.230 | -1.316 | .198 | | | | |
| **Step 5** | | | | - | .000 | .000 | -.053 |
| (Constant) | - | 8.264 | .000 | | | | |

*Note.* N = 33. Each row represents one step in the hierarchical regression analysis, i.e., after removing a non-significant predictor from the model. FO = fractional occupancy. *p < .05, ** p < .01.

**Supplementary Table 3.** General HP as dependent variable and mean duration values of states 1-5 as predictors.

| | Coefficients | | | Model summary | | | |
|---|---|---|---|---|---|---|---|
| **Predictor** | **β** | **t** | **p** | **F** | **$R^2$** | **Adj. $R^2$** | **$R^2\Delta$** |
| **Step 1** | | | | .771 | .125 | -.037 | .125 |
| (Constant) | - | .654 | .519 | | | | |
| State 1 MD | .058 | .285 | .778 | | | | |
| State 2 MD | -.311 | -1.478 | .151 | | | | |
| State 3 MD | .046 | .299 | .821 | | | | |

| | | | | | | | |
|---|---|---|---|---|---|---|---|
| State 4 MD | .279 | 1.414 | .169 | | | | |
| State 5 MD | .150 | .640 | .527 | | | | |
| **Step 2** | | | | .984 | .123 | -.002 | -.002 |
| (Constant) | - | .967 | .342 | | | | |
| State 1 MD | .058 | .288 | .775 | | | | |
| State 2 MD | .294 | -1.515 | .141 | | | | |
| State 4 MD | .268 | 1.424 | .165 | | | | |
| State 5 MD | .131 | .608 | .548 | | | | |
| **Step 3** | | | | 1.326 | .121 | .030 | -.003 |
| (Constant) | - | 1.067 | .295 | | | | |
| State 2 MD | -.304 | -1.613 | .118 | | | | |
| State 4 MD | -278 | 1.530 | .137 | | | | |
| State 5 MD | .160 | .853 | .401 | | | | |
| **Step 4** | | | | 1.641 | .099 | .039 | -.022 |
| (Constant) | - | 2.099 | .044 | | | | |
| State 2 MD | -.249 | -1.413 | .168 | | | | |
| State 4 MD | .243 | 1.378 | .178 | | | | |
| **Step 5** | | | | 1.344 | .042 | .011 | -.057 |
| (Constant) | - | 2.411 | .022 | | | | |
| State 2 MD | -.204 | -1.159 | .255 | | | | |
| **Step 6** | | | | - | .000 | .000 | -.042 |
| (Constant) | - | 8.264 | .000 | | | | |

*Note.* N = 33. Each row represents one step in the hierarchical regression analysis, i.e., after removing a non-significant predictor from the model. MD = mean duration. *p < .05, ** p < .01.

**Supplementary Table 4.** AH-HP as dependent variable and FO values of states 1-5 as predictors.

| | **Coefficients** | | | **Model summary** | | | |
|---|---|---|---|---|---|---|---|
| **Predictor** | **β** | **t** | **p** | **F** | **$R^2$** | **Adj. $R^2$** | **$R^2\Delta$** |
| **Step 1** | | | | 1.941 | .217 | -.105 | .217 |
| (Constant) | - | .513 | .612 | | | | |
| State 1 FO | .391 | -.086 | .704 | | | | |
| State 2 FO | -.086 | -.384 | .704 | | | | |
| State 3 FO | -.055 | -.252 | .803 | | | | |
| State 5 FO | -.085 | -.466 | .645 | | | | |
| **Step 2** | | | | 2.652 | .215 | .134 | -.002 |
| (Constant) | - | .510 | .614 | | | | |
| State 1 FO | .431 | 2.253* | .032* | | | | |
| State 2 FO | -.061 | -.309 | .760 | | | | |
| State 5 FO | -.087 | -.488 | .629 | | | | |
| **Step 3** | | | | 4.053 | .213 | .160 | -.003 |

| | | | | | | | |
|---|---|---|---|---|---|---|---|
| (Constant) | - | .438 | .664 | | | | |
| State 1 FO | .461 | 2.827** | .008** | | | | |
| State 5 FO | -.107 | -.656 | .517 | | | | |
| **Step 4** | | | | 7.818 | .201 | .176 | -.011 |
| (Constant) | - | -.066 | .948 | | | | |
| State 1 FO | .499 | 2.796** | .009** | | | | |

*Note.* N = 33. Each row represents one step in the hierarchical regression analysis, i.e., after removing a non-significant predictor from the model. FO = fractional occupancy. *p < .05, ** p < .01.

**Supplementary Table 5.** A-HP as dependent variable and mean duration values of states 1-5 as predictors.

| | **Coefficients** | | | **Model summary** | | | |
|---|---|---|---|---|---|---|---|
| **Predictor** | **β** | **t** | **p** | **F** | **$R^2$** | **Adj. $R^2$** | **$R^2\Delta$** |
| **Step 1** | | | | 1.502 | .218 | .078 | .218 |
| (Constant) | - | .627 | .536 | | | | |
| State 1 MD | .401 | 2.079* | .047* | | | | |
| State 2 MD | -.199 | -1.003 | .325 | | | | |
| State 3 MD | -.087 | -.454 | .654 | | | | |
| State 4 MD | .020 | .110 | .914 | | | | |
| State 5 MD | -.033 | -.150 | .882 | | | | |
| **Step 2** | | | | 1.943 | .217 | .105 | .000 |
| (Constant) | - | .718 | .479 | | | | |
| State 1 MD | .405 | 2.173* | .038* | | | | |
| State 2 MD | -.192 | -1.043 | .306 | | | | |
| State 3 MD | -.092 | -.502 | .619 | | | | |
| State 5 MD | -.042 | -.204 | .840 | | | | |
| **Step 3** | | | | 2.666 | .216 | .135 | -.001 |
| (Constant) | - | .723 | .476 | | | | |
| State 1 MD | .389 | 2.321* | .028* | | | | |
| State 2 MD | -.207 | -1.234 | .227 | | | | |
| State 3 MD | -.080 | -.470 | .642 | | | | |
| **Step 4** | | | | 3.991 | .210 | .158 | -.006 |
| (Constant) | - | .578 | .568 | | | | |
| State 1 MD | .405 | 2.497* | .018* | | | | |
| State 2 MD | -.222 | -1.367 | .182 | | | | |
| **Step 5** | | | | 5.949 | .161 | .134 | -.049 |
| (Constant) | - | -.920 | .365 | | | | |
| State 1 MD | .401 | 2.439* | .021* | | | | |

*Note.* N = 33. Each row represents one step in the hierarchical regression analysis, i.e., after removing a non-significant predictor from the model. MD = mean duration. *p < .05, ** p < .01.

**Supplementary Table 6.** AV-HP as dependent variable and FO values of states 1-5 as predictors.

| | Coefficients | | | Model summary | | | |
|---|---|---|---|---|---|---|---|
| **Predictor** | **β** | **t** | **p** | **F** | **$R^2$** | **Adj. $R^2$** | **$R^2\Delta$** |
| **Step 1** | | | | 1.616 | .188 | .072 | .188 |
| (Constant) | - | -.156 | .878 | | | | |
| State 1 FO | .470 | 1.841 | .076 | | | | |
| State 2 FO | .152 | .665 | .511 | | | | |
| State 3 FO | .001 | .006 | .995 | | | | |
| State 5 FO | -.228 | -1.233 | .288 | | | | |
| **Step 2** | | | | 2.232 | .188 | .104 | .000 |
| (Constant) | - | -.242 | .811 | | | | |
| State 1 FO | .469 | 2.406* | .023* | | | | |
| State 2 FO | .152 | .754 | .457 | | | | |
| State 5 FO | -.228 | -1.257 | .219 | | | | |
| **Step 3** | | | | 3.108 | .172 | .116 | -.016 |
| (Constant) | - | .563 | .578 | | | | |
| State 1 FO | .395 | 2.361* | .025* | | | | |
| State 5 FO | -.177 | -1.061 | .297 | | | | |
| **Step 4** | | | | 5.070 | .141 | .113 | -.031 |
| (Constant) | - | -.318 | .753 | | | | |
| State 1 FO | .375 | 2.52* | .032* | | | | |

*Note.* N = 33. Each row represents one step in the hierarchical regression analysis, i.e., after removing a non-significant predictor from the model. FO = fractional occupancy. *$p < .05$, ** $p < .01$.

**Supplementary Table 7.** AV-HP as dependent variable and mean duration values of states 1-5 as predictors.

| | Coefficients | | | Model summary | | | |
|---|---|---|---|---|---|---|---|
| **Predictor** | **β** | **t** | **p** | **F** | **$R^2$** | **Adj. $R^2$** | **$R^2\Delta$** |
| **Step 1** | | | | 1.041 | .162 | .006 | .162 |
| (Constant) | - | .161 | .874 | | | | |
| State 1 MD | .426 | 2.132* | .042* | | | | |
| State 2 MD | .010 | .049 | .962 | | | | |
| State 3 MD | -.090 | -.457 | .651 | | | | |
| State 4 MD | -.034 | -.176 | .861 | | | | |
| State 5 MD | -.145 | -.631 | .534 | | | | |
| **Step 2** | | | | 1.349 | .162 | .042 | .000 |
| (Constant) | - | 1.74 | .863 | | | | |
| State 1 MD | .424 | 2.191* | 0.037* | | | | |

| | | | | | | | |
|---|---|---|---|---|---|---|---|
| State 3 MD | -.087 | -.477 | .637 | | | | |
| State 4 MD | -.031 | -.173 | .864 | | | | |
| State 5 MD | -.140 | -.695 | .493 | | | | |
| **Step 3** | | | | 1.850 | .161 | .074 | -.001 |
| (Constant) | - | .114 | .919 | | | | |
| State 1 MD | .419 | 2.225* | .034* | | | | |
| State 3 MD | -.083 | -.465 | .645 | | | | |
| State 5 MD | -.132 | -.685 | .499 | | | | |
| **Step 4** | | | | 2.739 | .154 | .098 | -.006 |
| (Constant) | - | -.281 | .780 | | | | |
| State 1 MD | .427 | 2.304* | .028* | | | | |
| State 5 MD | -.112 | -.604 | .550 | | | | |
| **Step 5** | | | | 5.219 | .144 | .116 | -.010 |
| (Constant) | - | -1.178 | .248 | | | | |
| State 1 MD | .380 | 2.285* | .029* | | | | |

*Note.* N = 33. Each row represents one step in the hierarchical regression analysis, i.e., after removing a non-significant predictor from the model. MD = mean duration. *p < .05, ** p < .01.

### C. Additional results of the source localization

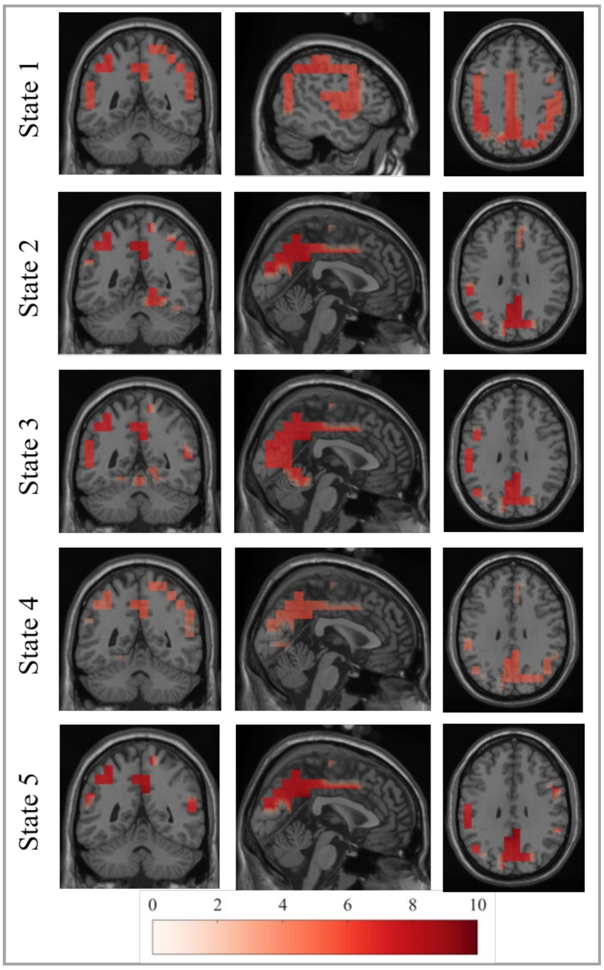


**Supplementary Fig. 1.** Orthogonal slices of highest mean source activation per state

The images show the cortical slices corresponding to the source of highest activation strength per state. Activation is expressed as Neural Activity Index (NAI; Van Veen et al. 1997). Images were generated by i) parcellating the individual source images using the AAL atlas into 116 areas, ii) averaging the activity within each parcel across subjects, iii) thresholding the mean sources images retaining only the highest 25% of active areas (>/= z-value of 0.675), and iv) interpolating the active sources onto a template MRI provided by Fieldtrip. State 1 showed the highest activation in the left inferior parietal lobe [MNI -50 -50 40]. State 2-5 showed the highest activation in the left posterior cingulate cortex [MNI 0 -50 30].

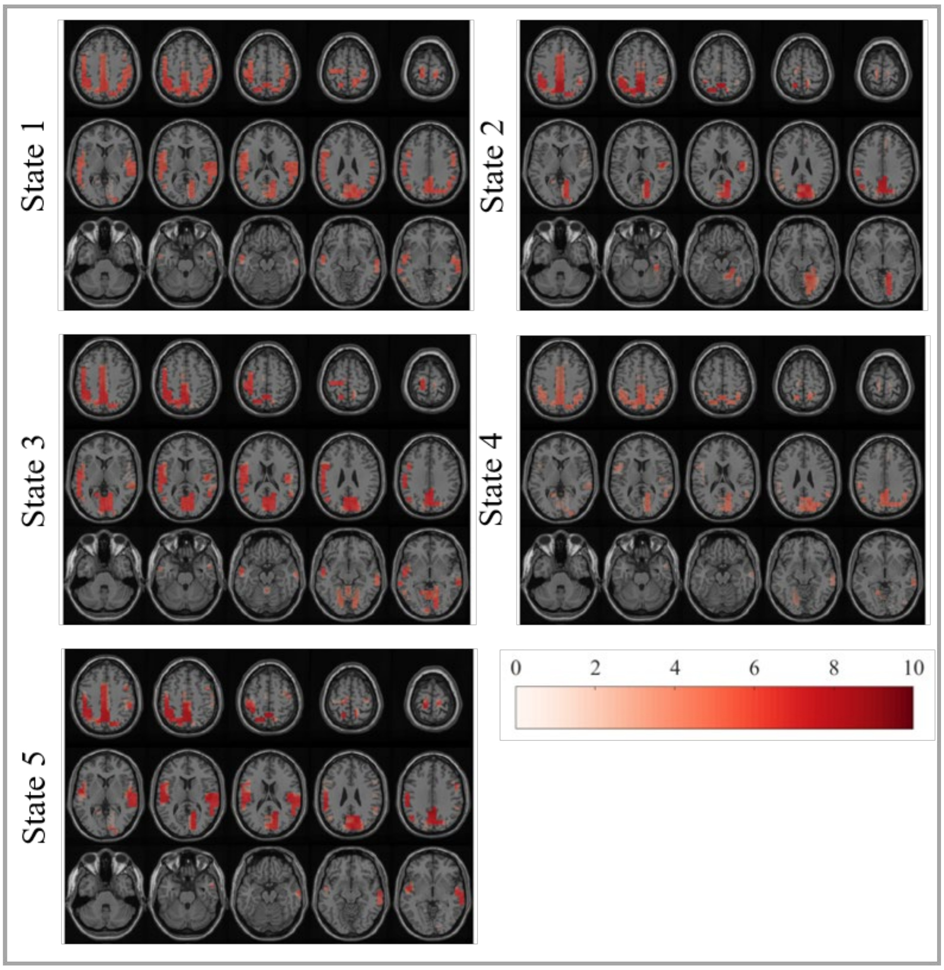


**Supplementary Fig. 2.** Axial slices of mean source activation per state

Slices show the source of highest activation strength per state. Activation is expressed as Neural Activity Index (NAI; Van Veen et al. 1997). Images were generated by i) parcellating the individual source images using the AAL atlas into 116 areas, ii) averaging the activity within each parcel across subjects, iii) thresholding the mean sources images retaining only the highest 25% of active areas (>/= z-value of 0.675), and iv) interpolating the active sources onto a template MRI provided by Fieldtrip.
